# Persistent enrichment of multidrug resistant *Klebsiella* in oral and nasal communities during long-term starvation

**DOI:** 10.1101/2023.12.18.572173

**Authors:** Jett Liu, Nell Spencer, Daniel R. Utter, Alex Grossman, Nídia C.D. Santos, Wenyuan Shi, Jonathon L. Baker, Hatice Hasturk, Xuesong He, Batbileg Bor

**Author notes:** These authors contributed equally (Jett Liu, Nell Spencer). Correspondence (B.B.).

## Abstract

The human oral and nasal cavities can act as reservoirs for opportunistic pathogens capable of causing acute infection. These microbes asymptomatically colonize the human oral and nasal cavities which facilitates transmission within human populations via the environment, and they routinely possess a clinically-significant antibiotic-resistance genes. Among these opportunistic pathogens, the *Klebsiella* genus stands out as a notable example, with its members frequently linked to nosocomial infections and multidrug resistance. As with many colonizing opportunistic pathogens, how *Klebsiella* transitions from an asymptomatic colonizer to a pathogen remains unclear. Here, we explored a possible explanation by investigating the ability of oral and nasal *Klebsiella* to outcompete their native microbial community members under *in vitro* starvation conditions, which could be analogous to external hospital environments. When *Klebsiella* was present within a healthy human oral or nasal sample, the bacterial community composition shifted dramatically under starvation conditions and typically became dominated by *Klebsiella*. Furthermore, introducing *K. pneumoniae* exogenously into a native microbial community lacking *K. pneumoniae*, even at low inoculum, led to repeated dominance under starvation. *K.pneumoniae* strains isolated from healthy individuals’ oral and nasal cavities also exhibited resistance to multiple classes of antibiotics and were genetically similar to clinical and gut isolates. In addition, we found that in the absence of *Klebsiella*, other understudied opportunistic pathogens, such as *Peptostreptococcus*, dominate under starvation conditions. Our findings establish an environmental circumstance that allows for the outgrowth of *Klebsiella* and other opportunistic pathogens. The ability to outcompete other commensal bacteria and to persist under harsh environmental conditions may contribute to the colonization-to-infection transition of these opportunistic pathogens.

## Introduction

Microbes that are typically of low abundance and harmless in their normal niche, but potentially dangerous under specific circumstances are termed colonizing opportunistic pathogens (COPs) [1]. Translocation of human oral bacteria to other body sites has been associated with noncommunicable, chronic systemic illnesses including diabetes, cancer, and Alzheimer’s disease [2–5]. The oral microbiota also contains potential opportunistic pathogens that can cause more acute diseases [6]. These oral COPs include pathogens such as *S. pyogenes, S. pneumoniae* and *H. influenzae,* which are part of the normal oral community but can cause numerous diseases, including but not limited to streptococcal pharyngitis, pneumonia, sepsis, and/or meningitis [7]. Crucially, oral microbes and COPs can asymptomatically colonize and transmit between persons, complicating their disease etiology [8]. Characterizing how oral COPs colonize, transmit, and transition from commensal to pathogen is crucial, given that human oral fluids can contaminate surfaces and be ingested, disseminating throughout the body [9–12].

*Klebsiella* are a prominent example of an oral COP. Typically, oral *Klebsiella* species exhibit a low prevalence and abundance in populations studied to date, [13–15] having an increased association with periodontal pockets [14,16,17]. Despite their relatively low prevalence in the oral microbiome, whenever they are found, *Klebsiella* seem to be a consistent colonizer and member of the community [15]. Oral fluids are a major source of contamination in the healthcare setting, particularly in sinks and drains [12,18], and previous studies have demonstrated that *Klebsiella* can persist and transmit particularly well in nosocomial environments [19,20]. Therefore, a small number of individuals carrying oral *Klebsiella* could potentially act as superspreaders, seeding infection within a larger population and in at-risk persons.

Although routinely carried asymptomatically, *Klebsiella* are one of the six ESKAPE pathogens that can be highly virulent and are routinely resistant to multiple antibiotics [21,22]. As a result, *Klebsiella* is a major cause of life-threatening hospital-acquired infections in at-risk immunocompromised and critically ill patients [23]. *Klebsiella* species are also associated with various gastrointestinal infections, which can lead to gastroenteritis, colitis, and other chronic diseases [24]. The origin of *Klebsiella* infection has been linked to environmental samples (including soils, plants, and animals), human body sites (including the gut and skin microbiomes), and hospital environments (including contaminated sinks and drains) [13,19–21,25,26]. Despite the strong linkages between *Klebsiella* and nosocomial infections, the role of the human oral cavity as a reservoir for *Klebsiella* associated with nosocomial and gut infections has been largely overlooked. Most research on oral *Klebsiella* as an opportunistic pathogen was conducted in mouse studies and has demonstrated that mouse or human oral *Klebsiella* species can migrate to the mouse gut and cause various inflammation and colitis [27,28].

Illustrating its toughness and ability to survive in various stress conditions, numerous nosocomial pathogens, including *Klebsiella*, have been shown to persist on various surfaces [29,30]. These bacteria may possess various mechanisms to withstand environmental stress, enabling them to survive challenging conditions for varying durations. While these adaptations may have evolved in response to environmental stress, many environmental stress responses are often useful for pathogenic niches, including desiccation resistance [31], chemical competition [32], and persistence states [33]. Our previous study examined the oral microbiome community under starvation conditions akin to those of an expelled oral droplet persisting on a surface [34]. We found that members of the Enterobacteriaceae family, and particularly *Klebsiella* species, outlasted other bacterial community members to emerge as the only surviving taxa. This finding was assessed using metagenomics, metatranscriptomics, and traditional growth methods, confirming that, among the studied samples, Enterobacteriaceae were the majority of the transcriptionally active bacteria after >100 days of starvation, and the only taxa that could be recovered by plating.

The previous study, however, was limited by a small sample size and did not examine the ubiquity of the Enterobacteriaceae survival phenomenon. In this study, we further assessed the capability of *Klebsiella* to persist under starvation conditions by including additional clinical samples from the oral cavity. We also examined nares samples given that nares fluids can contaminate hospital surfaces. We observed that when *Klebsiella* species were present within a community, they consistently dominated starved communities, especially *Klebsiella pneumoniae,* which not only endured starvation but also proliferated rapidly to establish dominance within the first 24 hours of starvation. Furthermore, all cultured strains of *Klebsiella pneumoniae* isolates from healthy human oral and nasal cavities exhibited multidrug resistance. These strains were found to be phylogenetically intermingled with previously sequenced clinically relevant and healthy gut isolates. In the absence of *Klebsiella* species, other bacteria such as *Peptostreptococcus* dominate starved communities. These findings collectively suggest that oral COPs, such as *Klebsiella* species, have the capability to persist longer than other members of oral and nasal microbial communities under starvation conditions. This enduring trait might have the potential to influence the colonization-to-infection process, especially in areas where the opportunistic pathogen has to go through starvation or other stress environments.

## Results

### Starvation of oral and nasal polymicrobial communities in vitro reveals enriched genera

Thirty healthy human volunteers were recruited for the sampling of two body sites, saliva (S1-S30) and nares (N1-N30), for a total of sixty samples. These initial oral and nasal communities (hereafter referred to as the raw communities) were inoculated into SHI-medium, and resultant communities were incubated for 24 hours (hereafter referred to as the SHI-medium communities). SHI-medium has been extensively used to expand the biomass of oral microbial communities with minimal impact on biodiversity [35]. Subsequently, the SHI-medium communities were resuspended and aliquoted in PBS and starved under aerobic conditions for both 30 days (hereafter referred to as the day-30 communities) and 120 days (hereafter referred to as the day-120 communities) (Figure 1A).

**Figure 1.**
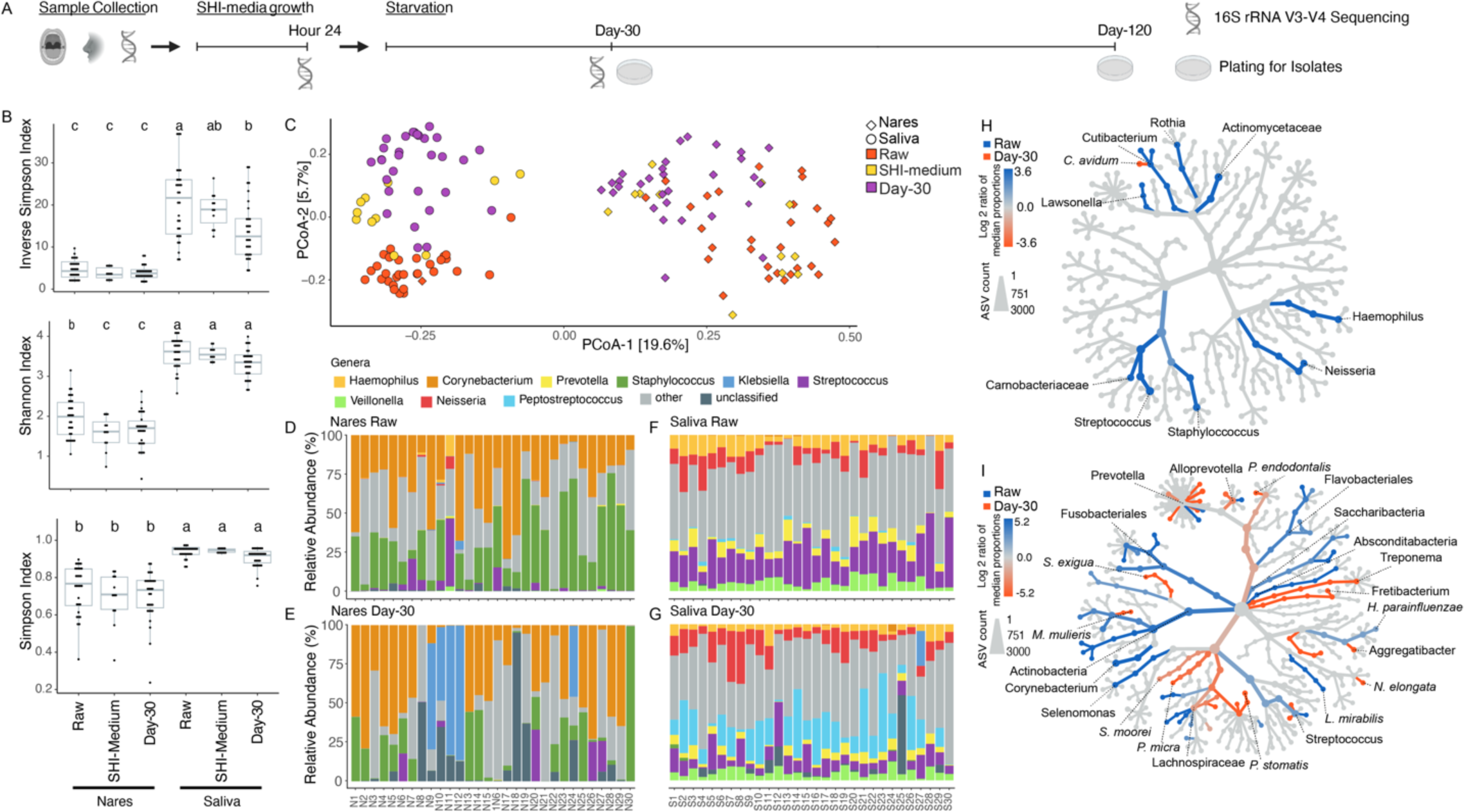
Community and relative abundance analysis of sequenced samples. A) Timeline detailing sample collection, SHI-medium growth, and starvation procedures. Sequencing and plating times are indicated. B) Alpha diversity analysis of sequenced samples grouped by culture condition and body site. Alpha diversity was calculated using ASV abundances. C) Principal coordinates analysis (PCoA) plot produced using the Bray-Curtis distance function and ASV abundances. Body site and culture condition are indicated by color. D-G) Stacking bar charts detailing the relative abundance of genera identified within sequenced samples. H) Comparative analysis of bacteria gained or lost between nares raw communities and day-30 communities. Color indicates lost or gained taxon groups on a log_2_ median proportion scale. Node size indicates normalized ASV counts. Only taxon levels with a Wilcoxon rank-sum p-value ≤ 0.05 are colored. I) Comparative analysis of bacteria gained or lost between saliva raw communities and day-30 communities. Color indicates lost or gained taxon groups on a log_2_ median proportion scale. Node size indicates normalized ASV counts. Only taxon levels with a Wilcoxon rank-sum p-value ≤ 0.05 are colored.

To initially assess how starvation conditions alter the composition of oral and nasal communities, we performed 16S rRNA gene sequencing of the V3-V4 region on all raw communities, several SHI-medium communities, and all day-30 communities. Using identified ASVs, we analyzed the alpha (intrasample) diversity of the different communities using the Shannon, Simpson, and Inverse Simpson indices. Broadly, nares samples were less diverse than saliva samples (Figure 1B), an expected finding that has been reported in other studies [36]. Across both nares and saliva samples, we observed a slight decrease in alpha diversity when comparing the raw communities to the SHI-medium communities (Figure 1B). Further, both nares and saliva day-30 communities tended to exhibit slightly lower alpha diversity when compared to the SHI-medium communities (Figure 1B). Altogether, the alpha diversity metrics generally indicated that both nares and saliva communities tended to decrease in alpha diversity when subjected to starvation conditions. Beta diversity (intercommunity) plots based on identified ASVs displayed a clear separation between raw nares and saliva communities (PERMANOVA, p ≤ 0.001), an expected result given the known difference in bacterial community composition across body sites (Figure 1C). Similarly, samples from the same body site are typically clustered according to culture condition (Figure 1C), an observation particularly pronounced among the nares samples. In general, SHI-medium communities had a less pronounced shift away from the raw communities when compared to day-30 communities, which clearly clustered apart from their raw community counterparts.

We next analyzed the genera-level relative abundance data, which revealed several interesting trends. Although the nares samples did not appear to display a consistent increase in a particular genus when comparing the raw communities to the day-30 communities, it was striking to observe that five of the thirty day-30 communities (N9, N10, N11, N12, and N24) contained a large relative abundance - greater than 45% - of *Klebsiella* (Figure 1D and 1E, Table S1, S2, and S3). This large increase in *Klebsiella* abundance was even more pronounced when considering that only six raw nares communities (Samples N9, N10, N11, N12, N17, N24) contained detectable levels of *Klebsiella,* with a mean relative abundance of ∼1.28% (0.1-5.7% range). In contrast to the nares samples, saliva communities displayed more consistent changes in genera-level relative abundances. In particular, we broadly noted that compared to raw saliva communities day-30 communities decreased in *Streptococcus* relative abundance and increased in *Peptostreptococcus* relative abundance (Figure 1F and 1G, Tables S1, S2, and S3). We observed only one saliva sample (S27) had detectable *Klebsiella* in the raw communities, and its abundance increased to ∼22.5% of the day-30 community (Figure 1G). Interestingly, we did not detect *Klebsiella* in both the oral and nasal cavity of the same individual in any of the samples.

Global comparative analysis of community composition confirmed these initial observations. Nares samples displayed few consistent statistically significant changes when subjected to starvation (Wilcox rank sum ≤ 0.05) (Figure 1H, Table S4). In contrast, saliva samples were much more consistent in statistically significant community composition changes between raw and starvation conditions (Figure 1I, Table S5).

### Opportunistic pathogens dominate starved communities

Particularly intrigued by the dominance of *Klebsiella* in several of the nares and one saliva day-30 communities, we sought to unbiasedly assess if specific species were enriched in the starved communities (Figure 2A, 2B). At the species level, three nares raw communities contained a single species with above 50% relative abundance (Figure 2A, Table S6). These initially dominant species were *Dolosigranulum pigrum* (N2 and N22) and *Corynebacterium tuberculostearicum* (N12). Strikingly, after starvation, nine nares day-30 communities contained a single species with over 50% relative abundance (Figure 2A, Tables S7 and S8). These nine dominant species were *Klebsiella pneumoniae*, *Klebsiella aerogenes*, *Kocuria rhizophila*, *Cronobacter sakazakii*, *Bacillus anthracis*, *Corynebacterium pseudodiphtheriticum*, *Anaerococcus* sp. HMT-295, *Staphylococcus warneri*, and *Granulicatella adiacens* (Figure 2C). Notably, many of these species, including *Klebsiella*, are known pathogens that have the ability to survive challenging thermal, desiccation and pH stresses environments [37–40]. Additionally, some of these species are not well known nasal commensals but have been detected sporadically [41]. This dominance of a particular species within a day-30 community was typically a single occurrence per taxa - 7 of the 9 dominant species had a relative abundance above 50% in only a single day-30 community (*K. rhizophila*, *C. skazakii, B. anthracis, C. pseudodiphtheriticum, A.* sp. HMT-295, *S. warneri*, and *G. adiacens*). Both *Klebsiella* species, *K. pneumoniae and K. aerogenes*, however, were present in multiple samples and displayed a consistent increase in relative abundance after starvation (Figure 2C).

**Figure 2.**
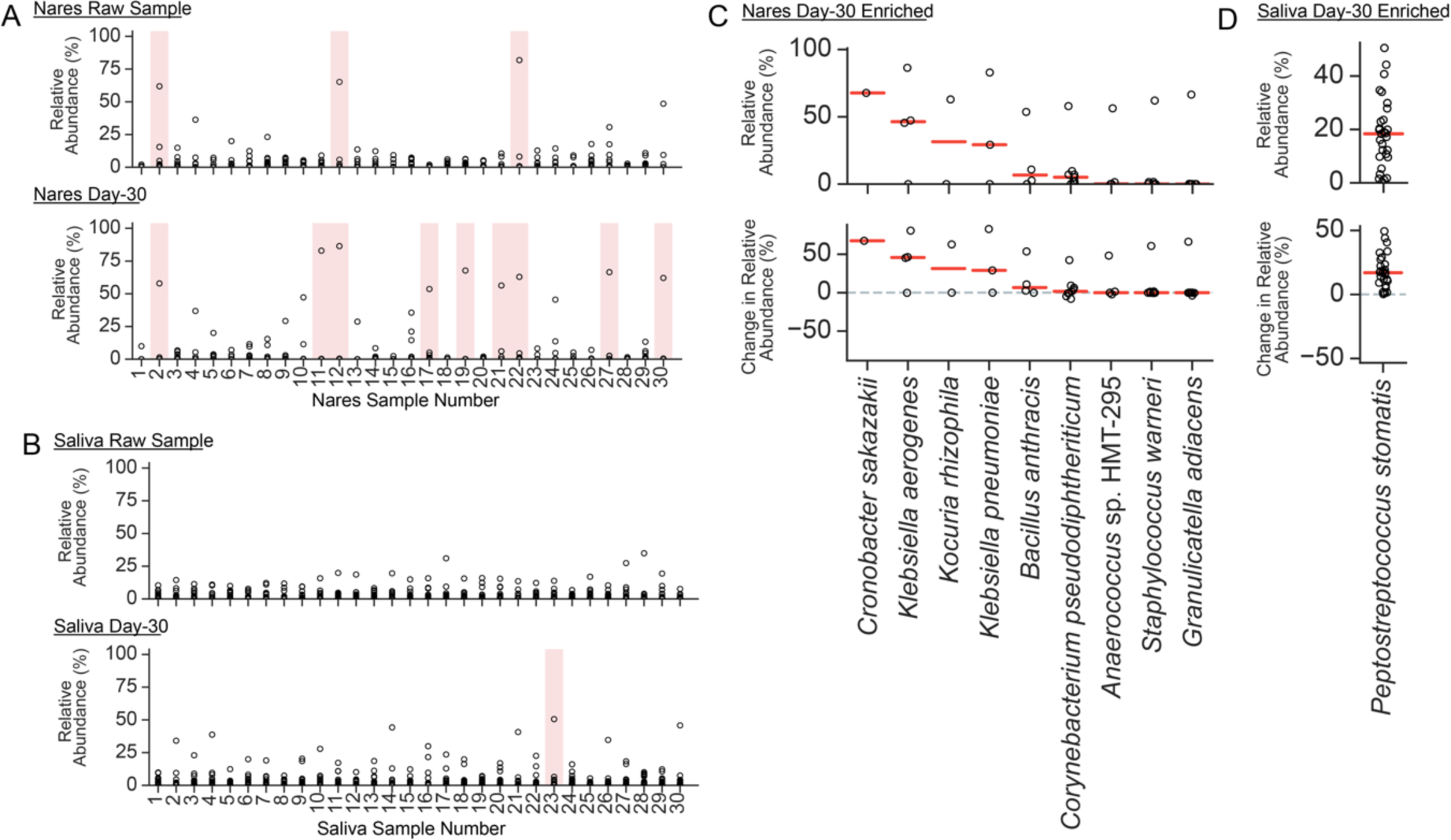
Species level analysis on enriched community members in stressed condition. A) Nares raw community and day-30 community species-based relative abundance analysis. Each point represents a single species. Samples (columns) with at least one species ≥ 50% relative abundance are shaded pink. B) Saliva raw community and day-30 community species-based relative abundance analysis. Each point represents a single species. Samples (columns) with at least one species ≥ 50% relative abundance are shaded pink. C) Relative abundance analysis of Nares species with ≥ 50% relative abundance in at least one day-30 community (shaded pink from panel A). Top panel displays day-30 relative abundances. Bottom panel displays the change in relative abundance from raw communities to day-30 communities. D) Relative abundance analysis of Saliva species with ≥ 50% relative abundance in at least one day-30 community (shaded pink from panel B). Top panel displays day-30 relative abundances. Bottom panel displays the change in relative abundance from raw communities to day-30 communities.

In contrast to the nares samples, the number of saliva samples with a dominant species above 50% relative abundance increased from zero in raw communities to one in day-30 community (Figure 2B, Tables S6, S7 and S8). This sample, day-30 S23, had a *P. stomatis* relative abundance greater than 50%. Additionally, across all samples, *P. stomatis* displayed a mean increase in relative abundance of 17.8% from raw to day-30 conditions (Figure 2D).

### Solid media growth of the starved communities largely support 16S rRNA sequencing results and dominated by Klebsiella species

16S rRNA gene sequencing detects both live and dead bacteria. To survey viable bacteria, we cultured communities from raw, day-30, and day-120 on blood BHI agar plates. Before starvation, raw communities exhibited a wide range of colony diversity; they varied broadly in size, color, and texture (Figure S1A). However, after 30 and 120 days of starvation, colony diversity decreased drastically, and we mostly observed uniform mucoid colonies (Figures 3, S1B-C). To identify the day-30 and day-120 colonies, we performed partial 16S amplification and Sanger sequencing on 91 colonies. In samples that had enriched *Klebsiella* by 16S rRNA profiling, the vast majority of the colonies from the starved communities were *Klebsiella* species (Figures 3A, 3B, Table S9). Specifically, two dominant *Klebsiella* species from independent samples emerged in our cultures, *K. pneumoniae* (samples N9, N11, and S27) and *K. aerogenes* (samples N10, and N24). When placed in a 16S rRNA phylogenetic tree, the sequenced *Klebsiella* formed two main clades accordingly (Figure 3G). Moreover, we identified and isolated multiple *Staphylococcus* species in both nares and saliva samples, *Rothia* from saliva samples, and *Cutibacterium* from nares samples (Figure 3C-F, Table S9). We did not observe growth of *Peptostreptococcus*, generally considered a strict anaerobe, suggesting the microaerophilic plating growth conditions did not support their growth [42]. Despite the limitations of blood BHI agar plates and microaerophilic conditions to support the growth of many saliva and nares bacteria, *Klebsiella* enrichment under starvation conditions, compared to raw communities, was very dramatic, suggesting that when present, even in small initial abundance, *Klebsiella* can take over the community and stay viable after starvation. In starvation cultures that did not have *Klebsiella*, we observed mixed colonies that were not dominated by one specific bacteria.

**Figure 3.**
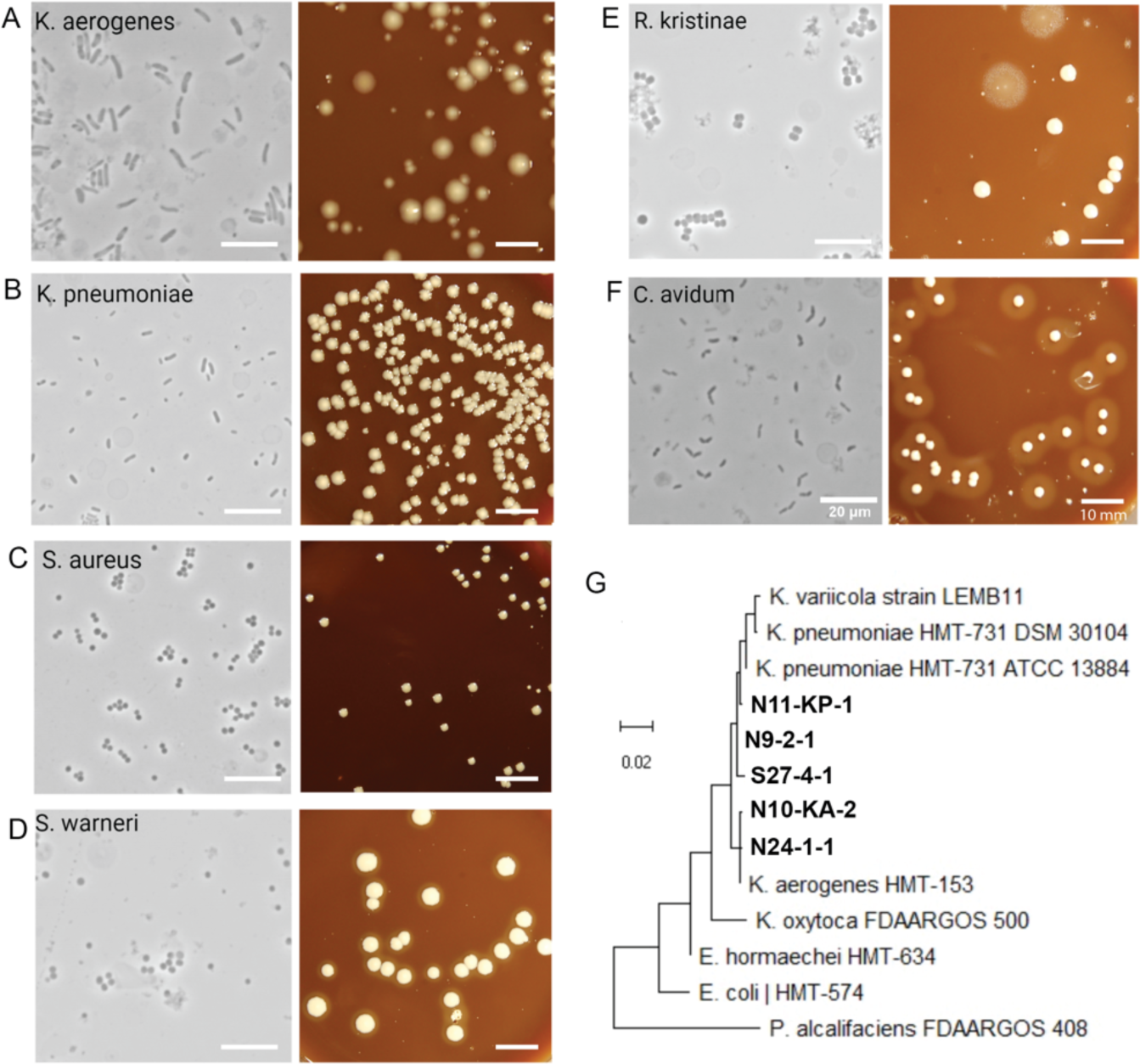
Growth of day-30 and day-120 communities on solid media. (A-F) Phase-contrast and colony images of isolated bacterial strains from the starved day-30 and day-120 communities. These are representative images of a larger cohort, which is summarized in Table S9. (G) Neighbor-joining tree for partial 16S rRNA sequences using MEGA X. Isolated bacterial strains from this study are designated by their nares (N) and saliva (S) subject numbers in bold and combined with representative reference strains from eHOMD. Only unique *Klebsiella* species from different saliva and nares samples were included. All scale bars are 20 micrometers for phase-contrast images, and 10 millimeters for colony images.

### Klebsiella pneumoniae dominates nares and saliva communities with high efficiency under starvation conditions

Considering that *Klebsiella* species were infrequently found in saliva and nares communities (1/30 samples and 5/30 samples), we aimed to strengthen the significance of our findings by conducting *in vitro* experiments. We randomly selected 15 saliva and 15 nares SHI-medium communities that lacked any presence of *Klebsiella* and spiked these samples with 0.1% relative abundance of *K. pneumoniae* strain N9-2-1 (Figure 4A). We picked N9-2-1 since, out of the three well-characterized *K. pneumoniae* strains in our study, it was the only strain that isolated from the day-120 starvation (Table S9). We reasoned that this strain would have more selective advantage compared to day-30 strains, and would show a clearer phenotype. The estimate of 0.1% was first determined from the 16S rRNA profiling of the raw sequence in Figure 1, and then optical density measurement and colony-forming units were used to estimate the final spiking (see methods). Post-spiking but before starvation, the *K. pneumoniae* relative abundance by 16S rRNA sequencing across the communities ranged from 0.04% to 21.0% (mean 4.3% relative abundance) (Figures 4B, 4C, 4D, Table S10).

**Figure 4.**
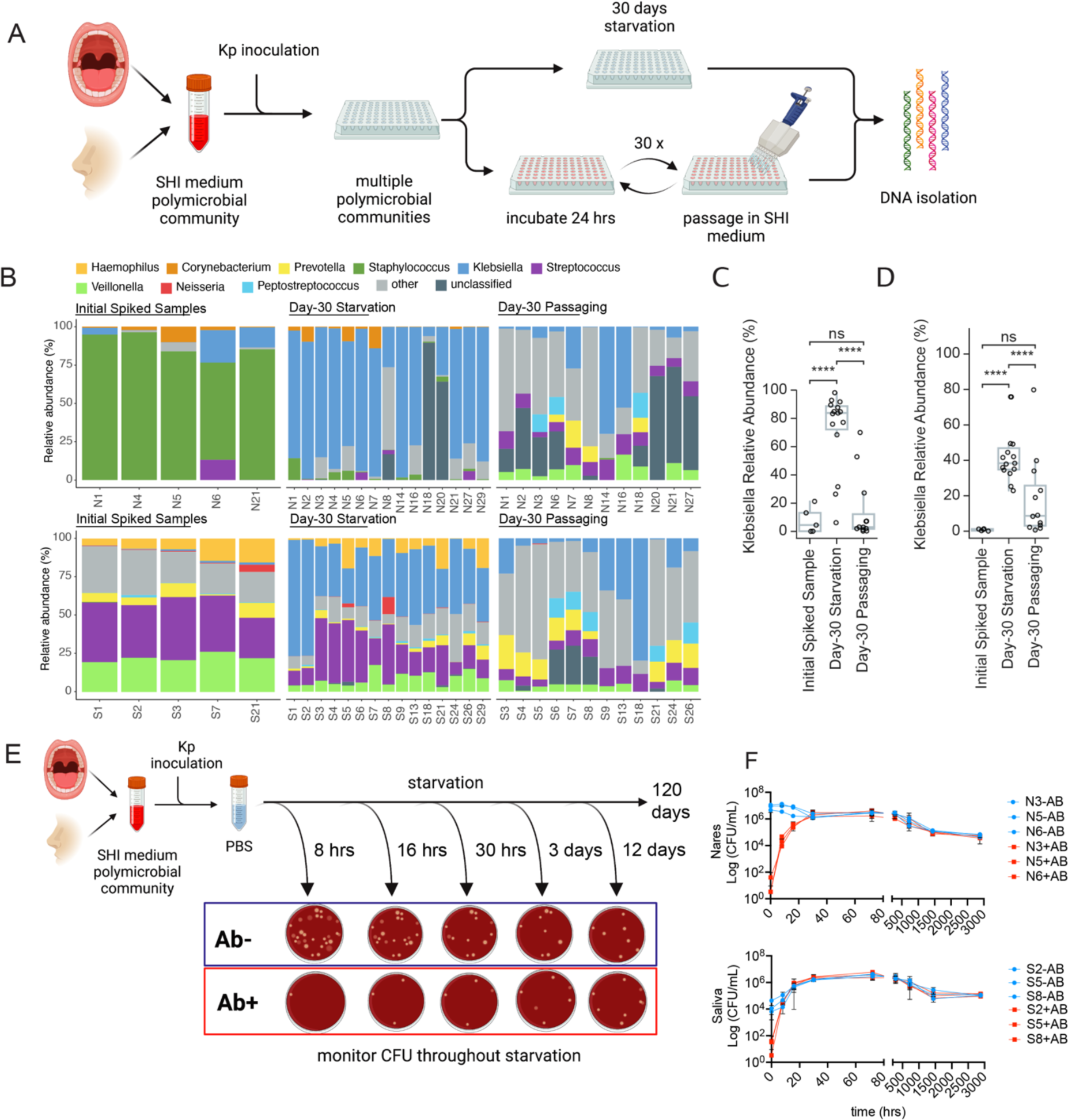
Artificial spiking of oral and nasal polymicrobial communities with *K. pneumoniae* result in consistent take over. (A) Cartoon depiction of *K. pneumoniae* spiking experiment. 15 saliva and 15 nares samples were spiked with low abundance of *K. pneumoniae* strains, and incubated by two different methods simultaneously: 30 days of starvation or 30 days of nutrition rich passaging in SHI media. (B) 16S relative abundance data of the spiking experiment after 30 days of starvation. Five samples of the initial spiked but not incubated community shown as a control. (C-D) Relative abundance of only *K. pneumoniae* plotted from panel B for all three groups. (E) Cartoon depiction of spiking experiment with determining viable colony forming units (CFU) of *K. pneumoniae* and other community members at each time point with (+AB) or without (-AB) antibiotic challenge. (F) CFUs were plotted against time of plating. Both CFU on BHI blood plate (blue, -AB) and antibiotic cocktail plate (red, +AB) are shown for three different SHI media communities of saliva and nares samples. We expect only antibiotic resistant *K. pneumoniae* will grow on +AB plates.

Subsequently, we subjected each spiked community to two conditions: one aliquot was starved for 30 days, while another was passaged in fresh SHI-medium at a 1:5 dilution every 24 hours for 30 days. Afterward, we analyzed the resulting communities by 16S rRNA gene sequencing. Strikingly, similar to the above starvation experiments, exogenously added *K. pneumoniae* expanded in almost all starved saliva and nares communities. Spiked and starved saliva communities contained a mean *K. pneumoniae* relative abundance of 42.7% (range: 22.8%-75.8%), and spiked nares communities contained a mean *K. pneumoniae* relative abundance of 72.2% (range: 6.2%-98.2%) (Figure 4B, C, D, Table S11). In contrast, the spiked communities that were passaged daily using fresh SHI-medium did not experience *K. pneumoniae* outgrowth to the same extent. Passaged spiked saliva communities contained a mean *K. pneumoniae* relative abundance of 18.8% (range 0.66%-79.8%), and passaged spiked nares communities contained a mean *K. pneumoniae* relative abundance of 14.7% (range 0.25%-70.0%) (Figure 4B, 4C, 4D, Table S12). We want to note that the SHI-medium passaging cohort did not experience a completely ideal nutrition-rich condition, as these cultures went through lag, exponential, and brief stationary phases during each passage. Nevertheless, despite these limitations, the results from both conditions clearly indicated that *K. pneumoniae* does not consistently dominate saliva or nares communities under nutrition-rich conditions. These 16S data were also supported by denaturing gradient gel electrophoresis (DGGE) band intensity analysis of communities N6, N21, S3 and S27 (Figure S2A).

Having obtained *K. pneumoniae* isolates from the healthy human oral cavity, we capitalized on the opportunity to experimentally characterize the antibiotic resistance of our isolated *Klebsiella.* Previous studies have categorized *Klebsiella pneumoniae* as a member of the ESKAPE pathogens, a group of antibiotic-resistant bacteria with high clinical relevance [21]. Initially, we evaluated the antibiotic and antibiotic cocktail resistance of *K. pneumoniae* strains isolated from three healthy individuals (N9-2-1, N11-KP-1, S27-4-1) (Table S9). We compared these *K. pneumoniae* strains to resistance of three SHI-media saliva and nares communities that did not contain any *Klebsiella* species. As expected, the *K. pneumoniae* strains demonstrated resistance to ampicillin, vancomycin, and the antibiotic cocktail comprising vancomycin, trimethoprim, cefsulodin, and amphotericin B (Figure S2B). Additionally, two strains (N9-2-1 and N11-KP-1) displayed partial resistance to Nalidixic acid, and a single strain (N9-2-1) displayed resistance to Erythromycin. Surprisingly, the three nares (N3, N5, N6) and three saliva (S2, S5, S8) SHI-medium communities were largely susceptible to all antibiotics that we tested.

Leveraging the antibiotic resistance of *Klebsiella* and the antibiotic susceptibility of the nares and saliva communities, we conducted another spiking experiment to further confirm that *Klebsiella* were dominating starvation communities when present. We again inoculated saliva (S2, S5, S8) and nares (N3, N5, N6) communities with *Klebsiella* before subjecting the spiked communities up to 120 days of starvation (Figure 4E). At several time points during the 120-day starvation, we plated each spiked community onto two types of blood agar plates: one that contained no antibiotics and one that contained the four-antibiotic cocktail of Vancomycin, Trimethoprim, Cefsulodin, and Amphotericin B. We determined the communities’ colony forming units (CFU) on each plate, which allowed us to assess the approximate relative abundance of viable *Klebsiella* in the communities during starvation. At the very early stages of starvation, we observed the presence of diverse colony morphology types distinct from the typical mucoid *K. pneumoniae* colonies on the antibiotic negative plates (Figure S3). Throughout the starvation period, *K. pneumoniae* initially displayed a low CFU count but quickly grew to take over the communities, within ∼30 hours and ∼8 hours for nares and saliva, respectively (Figure 4F). It is worth noting that, in saliva communities, *K. pneumoniae* grew to an even higher CFU count than the starting CFU count of the community, suggesting that either *K. pneumoniae* has a remarkable capability to take over in starvation, or large portion of the saliva community members such as *Peptostreptococcus* could not be cultured on blood BHI agar plates (Figure 4F, S3). *K. pneumoniae* managed to maintain a high presence throughout the entire starvation period. Starved day-60 and day-120 communities still contained ∼10^6 CFUs, and the majority of them were mucoid colonies. These findings reaffirm that *K. pneumoniae* has the potent ability to outcompete other species and thrive under starvation conditions for an extended period of time.

### Pangenome analysis of K. pneumoniae isolates reveal conserved pathogenic and antibiotic resistant genes compared to clinical isolates

Among our cultured *K. pneumoniae* isolates, we performed whole genome sequencing on three strains (N9-2-1, N11-KP-1 and S27-4-1), each from a different individual, for pangenomic comparison (Table S9). Our aim was twofold: firstly, to compare our isolated, starvation-resistant *K. pneumoniae* to previously sequenced clinically relevant isolates, and secondly, to determine whether these strains encoded genomic characteristics that might account for their survival. From whole genome comparisons, we first noticed that one of our strains designated as *K. pneumoniae* by 16S rRNA gene sequencing, strain N11-KP-1 (Figure 3G), was in fact *K. variicola,* a closely related species to *K. pneumoniae* that can be pathogenic [21]. We next investigated whether isolates from healthy saliva or nares samples were phylogenetically distinct from disease-associated *K. pneumoniae*. To this end, we constructed a phylogenomic bacterial marker gene tree using publicly available *K. pneumoniae* genomes from a diverse array of environments (Figure 5A). This analysis indicated that our saliva and nares strains from healthy donors were most closely related to strains isolated from infected patients and hospital environments (Figure 5A). Further, strains isolated from oral and nasal sites were intermixed with previously sequenced strains isolated from the gut, skin, and systemic infections. Thus, it does not appear that the colonization of specific body sites or pathogenicity is confined to particular clades of *K. pneumoniae*.

**Figure 5.**
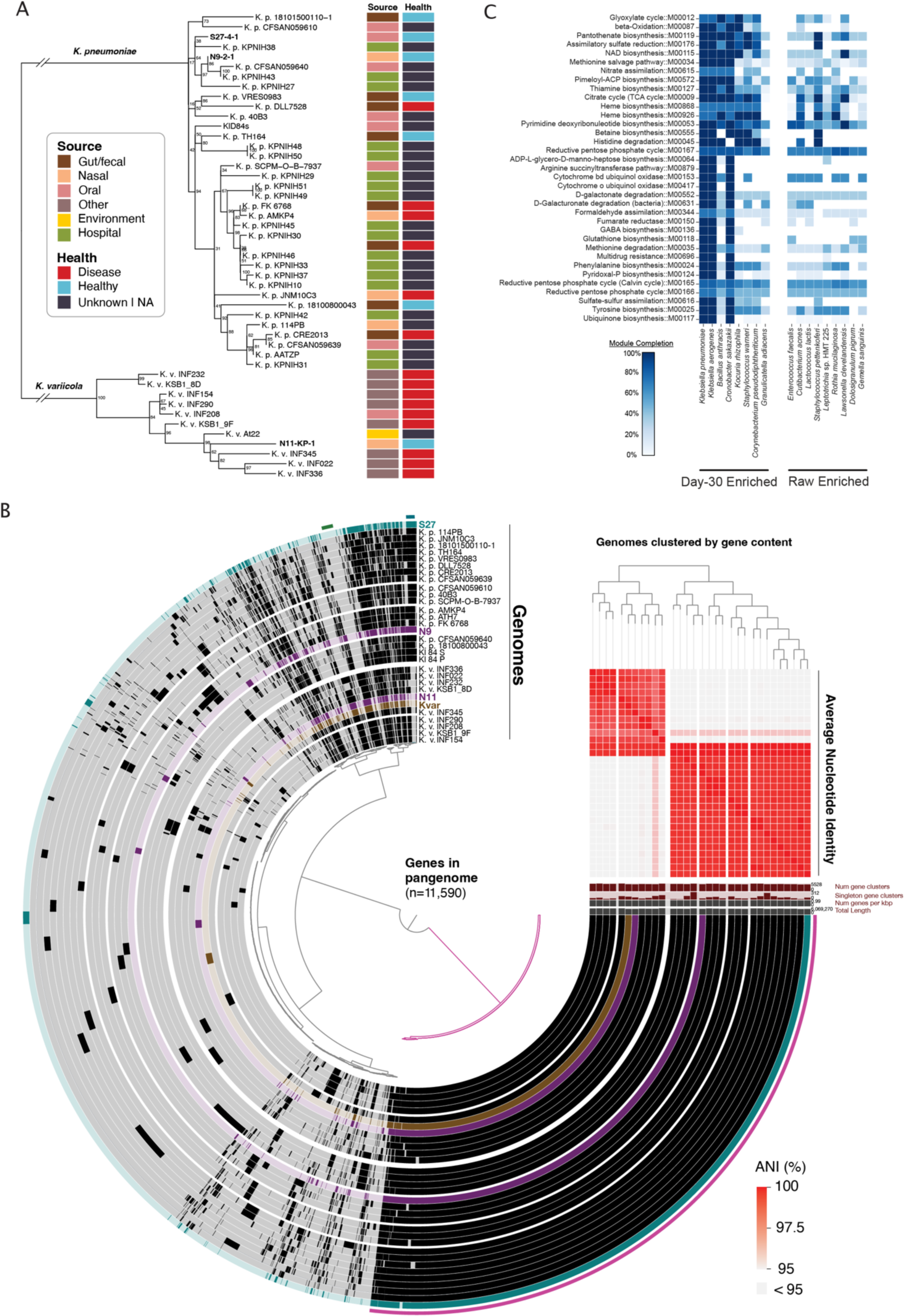
Genomic analysis of *K. pneumoniae* and *K. variicola*. (A) Maximum-likelihood tree of *Klebsiella* isolates from this study along with public reference genomes. Node labels represent bootstraps. Color boxes next to tips show the strain’s isolation source (first column) and the health status of the host (second column). (B) Pangenome of select *K. pneumoniae* and *K. variicola* isolates. Each track represents a different genome, and each unit along the track (i.e., inner dendrogram) represents a different gene homolog. Reference genomes are colored black, N9-2-1 and N11-KP-1 are purple, S27-4-1 is teal, and At22 (ant symbiont) is brown. Genomes are arranged by the presence/absence of genes (upper right dendrogram). The heatmap shows average nucleotide identity (ANI) between strains, clipped 95-100%. (C) Whole genome functional enrichment analysis of starvation enriched species compared to starvation depleted species. KEGG modules enriched among starvation enriched genomes with a p-value of < 0.05 are displayed. Color intensity signifies module completeness.

By constructing a pangenome to depict gene homolog presence or absence across genomes and clustering genomes based on the resultant data, the N9-2-1, N11-KP-1, and S27-4-1 isolates interspersed with other genomes from disparate isolation sources and host health statuses (Figure 5B). Notably, N9-2-1 and S27-4-1, our two *K. pneumoniae* isolates, shared more of their accessory genomes with other clinical isolates than with each other, reflected both in the visual display and the dendrogram. N11-KP-1 was more similar in gene content to the *K. variicola* At-22 isolate from a leafcutter ant colony than to most previously described *K. variicola* included here, although the two fell within and were overall highly similar to a group of clinical isolates. All genomes, including our new genomes and the re-analyzed reference genomes, also encoded multiple antibiotic resistance pathways (Figure S4A), consistent with our antibiotic testing in Figure S2B. The tool AMRFinder predicted resistance to ß-lactams, fosfomycin, phenicol/quinolone, tetracycline, and other efflux pumps important for antibiotic resistance (Table S13).

We also compared the genomes of the three isolates to a previously conducted saliva starvation experiment where *K. pneumoniae* genome mutants were tracked over 84 days [34]. Interestingly, all three of our isolates, sequenced after starvation, harbored the same nonsynonymous mutations found in Baker et al. 2019 which possibly gave conferred advantages to their survival (Figure S4B). Moreover, all other reference *K. pneumoniae* and *K. variicola* analyzed here (Figure S4B) shared the same or similar nonsynonymous mutations, suggesting that the majority of both clinical and carriage strains share preadaptation for stress or else these mutations are selected for during routine isolation.

Overall, the *Klebsiella* genomes displayed a high variability in gene content, perhaps reflecting their opportunistic lifestyle (Figure 5B). Yet, functionally, most genomes encode similar content, including similarities in nutrient and metal uptake, antibiotic resistance pathways, and more (Figure S5A). This observation is supported by previous reports of *Klebsiella* having large and highly dynamic genomes with a rich mobilome including plasmids, integrative conjugative elements, and other mobile elements [21]. Indeed, our assemblies found plasmids and mobile elements encoding genes for iron uptake (siderophores), antibiotic resistance, and more genes potentially useful to outlast and survive in harsh conditions (Table S14).

Given the high variability we observed among *K. pneumoniae* genomes and previous hypotheses that *Klebsiella* may be starvation-resistant due to their large genome size [34], we were curious if genome size could also explain why other species were also able to dominate occasionally under starvation conditions (Figure 2). To this end, we classified nares species present in our samples into two groups: 1) day-30 starvation enriched - those that were able to achieve a relative abundance greater than 50% in any sample (Figures 2A and 2D); and 2) day-30 starvation depleted - those whose raw community mean relative abundance decreased from greater than 1% to less than 0.1% mean relative abundance within the day-30 communities. Indeed, four of the eight day-30 starvation enriched species had a larger genome size than any raw enriched species: *K. pneumoniae*, *K. aerogenes*, *B. anthracis*, *C. sakazakii* (Figure S5B). Four post starvation enriched species, however, contained a genome with a similar size to the raw enriched species: *K. rhizophila*, *S. warneri*, *C. pseudodiphtheriticum*, *G. adiacens,* possibly indicating that genome size may not dictate starvation survivability in these species, rather specific gene or pathway is important.

We next performed functional enrichment analysis using whole genomes from eHOMD to assess if starvation-enriched species encoded similar metabolic pathways. Such a functional approach offers a means to disentangle genome size and functional potential, as genome size is also correlated with phylogeny and the majority of starvation-enriched isolates were Enterobacteraceae species. This analysis pointed towards several metabolic pathways that may be correlated with starvation survivability, many of which are involved in biosynthesis (Figure 5C). These genomes uniquely encoded several essential and non-essential amino acid biosynthesis and salvage pathways, including methionine, tyrosine, phenylalanine, arginine, histidine and thiamine, as well as carbohydrate metabolism functions, including D-galactonate degradation, succinate dehydrogenase, glyoxylate cycle and pentose phosphate cycle (Figure 5C).

## Discussion

Exploring the ecology and adaptations of starvation-resistant bacteria may provide insight into the infection mechanism of nosocomial pathogens since starvation conditions mimic the conditions endured within an expelled oral or nasal droplet residing on a hospital surface. Numerous studies have shown that *Klebsiella* and other opportunistic pathogens are hardy bacteria that can survive for long periods of time on different environmental surfaces [29,30], including hospital surfaces, sink drains, and ventilation systems [19,20,43,44]. The present study suggests that the oral and nasal cavities may act as a reservoir for colonizing opportunistic pathogens (COPs) such as *Klebsiella* and examined the ability of *Klebsiella* to dominate bacterial communities in starvation conditions. We demonstrated that *Klebsiella* species were exceptional and consistent in their ability to outlast and outgrow other members of the oral and nasal microbiomes under starvation conditions. Furthermore, these nasal and oral *Klebsiella* strains from healthy hosts were closely related to clinical isolates. They shared the majority of their gene clusters, including multidrug resistance genes, pathogenicity genes, and mobile elements, with strains isolated from nosocomial infections. Compared to species that were unable to survive starvation conditions, *Klebsiella pneumoniae* and other starvation-enriched species shared several metabolic pathways involved in biosynthesis that were not encoded among bacteria unable to survive starvation conditions. Interestingly, in the absence of *Klebsiella*, we saw saliva *Peptostreptoccocus* consistently increase in relative abundance upon starvation. Much like *Klebsiella*, previous literature has implicated *Peptostreptococcus* as a pathogen involved in hospital-acquired infections and periodontal disease [42,45,46].

*Klebsiella* is one of the ESKAPE pathogens and exhibits high virulence and resistance to antibiotics [21]. Our isolated *K. pneumoniae* strains possessed numerous antibiotic-resistant genes similar to clinical isolates. These strains also demonstrated natural resistance to more than three classes of antibiotics, satisfying the criteria for multidrug resistance (MDR) [47]. In comparison, the surrounding oral and nasal microbial community members, hundreds of species, were largely susceptible to the same antibiotics and antibiotic cocktails. In our study, we harnessed a distinctive resistance profile to selectively monitor the growth of *K. pneumoniae* in mixed communities. The natural occurrence of MDR strains within these polymicrobial communities presents a potential array of risks. These strains have the capacity to transfer their antibiotic resistance genes horizontally to other oral and nasal commensals or even to other pathogens passing through the oral cavity via our food sources [48,49]. Once they acquire antibiotic resistance in the oral and nasal cavities, these commensals, potential pathogens, or the MDR *K. pneumoniae* strains themselves, have multiple avenues to access vulnerable infection sites, including surface contamination or saliva swallowing. Consequently, antibiotic treatments may inadvertently amplify these organisms in the microbiome or infection, intensifying the level of risk [50,51]. Therefore, it is imperative to document and be cognizant of the presence of potentially threatening MDR oral commensal organisms.

How stable *Klebsiella* populations are within the oral or nasal cavity remains an open question. Given *Klebseilla*’s capability to survive and thrive in external environments and under harsh conditions, as demonstrated in this study, it is not illogical to consider that even a small number of patients or hospital staff harboring *Klebsiella* in their oral or nasal communities could inadvertently become key sources of *Klebsiella* contamination across broad settings. This is further amplified by the fact that *Klebsiella* can quickly outgrow other starving community members in our experiments and by 120 days of starvation, we still see large number of viable bacteria. Supporting the possibility that oral and nasal *K. pneumoniae* can disseminate, the whole genome comparison showed that the strains obtained here are essentially similar to other environmental and clinic isolates. We do not see taxonomic clustering of *K. pneumoniae* strains from particular body sites or infection, and there is no study to date suggesting that oral and nasal *K. pneumoniae* are essentially different from gut *K. pneumoniae*. With the current ability to sequence and assemble MAGs from different microbiome communities and the hospital environment, it is essential in the future to track and compare oral and nasal *K. pneumoniae* directly to strains from these environmental niches.

A better understanding of why the oral and nasal cavities typically exhibit a low prevalence and abundance of *Klebsiella* may elucidate novel ways to combat *Klebsiella* nosocomial infections. Previous analyses of the human microbiome project dataset showed that ∼9% of nasal samples contain *Klebsiella*, while only ∼3% of saliva samples contain *Klebsiella* [13]. *Klebsiella* abundance in these samples varies but is typically <1%. Our sampling of healthy oral and nasal cavities exhibited similar *Klebsiella* detection frequencies. 6/30 (20%) nares communities and 1/30 (3.3%) saliva samples were positive for *Klebsiella*. *Klebsiella* colonization of human body sites can depend on multiple factors such as the ability to bind epithelial cells, evasion of immune cells, competition for resources and space with native bacteria, and direct antagonism by native bacteria. In our study, in the absence of the eukaryotic hosts and in nutrient-rich conditions, *K. pneumoniae* was unable to outgrow native community members to the same extent as the starvation condition. Further, saliva communities, which harbored much more microbial diversity than nares communities, resisted *Klebsiella* outgrowth during starvation better than nares communities. These results suggest that there may be commensal saliva bacteria that modulate *Klebsiella* growth, either via nutrient competition or direct antagonism [52]. Consequently, there is a growing imperative for additional research on the ecology of *Klebsiella* and its communities. Such research holds the potential to offer insights into controlling *Klebsiella* and possibly other similar colonizing opportunistic pathogens.

In all, *Klebsiella* have the potential to dominate during starving, stressed conditions and this may give extra advantage outside the human oral and nasal cavity. Future studies will need to address how this exactly happens at the molecular level, and whether this contributes to the dissemination of these organisms to other body and infection sites.

## Acknowledgments

We thank Dr. Jeffery McLean, Dr. Deepak Chouhan, Susan Yost, and Lujia Cen for their productive discussion and material handling. This research was partially supported by grants from the National Institute of Dental and Craniofacial Research of the National Institutes of Health under Awards 1K99DE027719 (B.B.); and 1R01DE023810 (X.H.).

## Materials and Methods

### Clinical sampling

We collected 1.5 mL of unstimulated saliva samples (named S1-S30) and anterior nares swabs (named N1-N30) from a group of 30 adults comprised of 18 females and 12 males, all of whom had well-documented medical, dental, and periodontal health records. We excluded participants who had received periodontal disease treatment within the past 3 months, antibiotic treatment within the last 8 weeks, or antimicrobial mouthwash treatment within the last 4 weeks. Additionally, individuals with fewer than ten teeth or those currently using dentures were excluded from the study. The study received approval from the Forsyth Institutional Review Board under protocol number #18-06, and all subjects willingly provided written informed consent, demonstrating their voluntary participation in the research. To collect samples, unstimulated saliva was passively drooled into 15 mL conical tubes for collection, while moisten rayon-tipped swabs (Fisher Scientific #22-029-570) were inserted 1 cm into the anterior nares (separate for each nostril) and slowly rolled 5 times. Both swabs from the same individual were inserted into 2 mL sterile buffer saline and mixed vigorously to collect the microbiome.

### Polymicrobial community starvation

An aliquot of raw collected saliva and nares samples from each volunteer were reserved for genomic DNA isolation. The remaining fresh, non-frozen, saliva and nares samples were grown 24 hours in SHI-medium [35] in microaerophilic conditions (2% O2, 5% CO2, 93% N2) at 37°C. A portion of the SHI-medium culture was reserved for genomic DNA isolation. The remaining culture was centrifuged at 18,000 x g for 10 minutes and the pellet was resuspended in sterile phosphate buffer saline (PBS). Bacterial suspension in PBS derived from all 30 oral and 30 nares samples were further incubated at 37°C for 30 and 120 days with room air gas conditions with tubes sealed shut (hereafter referred to as the day-30 and day-120 communities). Following long-term starvation, a portion of these starved samples were reserved for genomic DNA isolation. The remaining samples were grown overnight in fresh SHI-medium at microaerophilic conditions at 37°C. The recovered bacteria were plated on blood agar media and colonies were isolated and identified via Sanger 16S rRNA gene sequencing (Table S9). All day-30 starvation communities were sequenced. Both day-30 and day-120 starvation communities were plated for bacterial isolation in microaerophilic gas environment.

### Genomic DNA isolation and Illumina sequencing

We isolated genomic DNA from saliva and nares samples at three time points: samples freshly collected from healthy subjects (raw communities), *in vitro* communities after expansion in SHI-medium prior to starvation (SHI-medium communities), and communities that were starved in PBS for 30 days (day-30 communities). We did not sequence day-120 communities because our previous study showed that the day-30 time point was adequate for showing the majority of the enriched species [28]. However, both day-30 and day-120 communities were grown on agar plate to determine viable bacteria and isolate enriched species (see above). Genomic DNA was isolated using the MasterPure Gram-Positive DNA purification Kit (Biosearch Technologies) with a modified protocol that included a bead-beating step [53]. Generally, gram-positive bacteria are harder to lyse, and thus, the modified kit was used to isolate both gram-negative and -positive bacterial genomic DNA. Briefly, cells were lysed using a lysis buffer and a cell disruption machine with beads. Proteinase K and a protein precipitation reagent were added to digest and allow for the removal of protein from the sample. Genomic DNA was precipitated in isopropanol and rinsed in ethanol. Extracted DNA was stored at -80°C until sequenced. Extracted DNA was sequenced by the Zymogen sequencing core on the Miseq Illumina sequencing platform targeting the V3-V4 region of the 16S rRNA gene.

### 16S RNA population profiling and analysis

16S rRNA ASVs were generated by a Zymogen in-house workflow that utilizes DADA2 [54]. ASV counts were normalized across samples using the Metacoder function “calc_obs_props” [55]. Using Metacoder, Shannon, Simpson, and Inverse Simpson alpha diversity metrics were calculated for each sample. PCoA beta diversity plots were generated using the Metacoder command “ordinate” and the specific flags “method = “PCoA”, distance = “bray””.

To assign taxonomy to the ASVs, we generated a custom Decipher classifier [56] using the most recent version of the extended Human Oral Microbiome Database (eHOMD) [41]: “https://www.google.com/url?q=https://www.homd.org/ftp/16S_rRNA_refseq/HOMD_16S_rRNA_RefSeq/V15.23/HOMD_16S_rRNA_RefSeq_V15.23.p9.fasta&sa=D&source=docs&ust=1693239952469670&usg=AOvVaw0M5dl1Ifonaz0Slth7IZWU”. To generate this custom classifier, the max group size was set to 10, and the max number of iterations were set to 3. We then used the Decipher IDTAXA approach [57] with the custom eHOMD database to assign taxonomy to the ASVs (Tables S1, S2, S3, S6, S7, S8). Taxon-level bar charts were generated using Phyloseq [58].

To compare the community member differences between the raw samples and starved samples, we generated differential heat trees using the Metacoder “heat_tree” function (Table S4, S5). Only differences with a Wilcox p-value less than 0.05 were displayed on the trees (“false discovery rate”). To display the trees, we used the “davidson-harel” layout algorithm.

### Genomic comparisons between nares bacterial species enriched and depleted under starvation conditions

We divided the species present within our nares communities into one of three groups: 1) starvation enriched - species that achieved 50% or greater relative abundance in any starved community; 2) starvation depleted - species with a mean relative abundance of 1% or greater in raw communities and a mean relative abundance of 0.1% or less in starved communities; 3) neutral - species that did not match the criteria for groups 1 or 2.

To assess if there was a correlation between genome size and starvation-enriched bacterial species, we used the known genome sizes listed on eHOMD. To compare the genomic inventories of starvation-enriched bacterial species to starvation-depleted bacterial species, we downloaded whole genomes of these bacterial species, when available, from eHOMD. Using anvi’o [59], we used the ‘anvi-estimate-metabolism’ function, which employs the KEGG module database [60], to annotate each genome. We then used the anvi’o ‘anvi-compute-metabolic-enrichment’ function at the default completeness threshold of 75% to identify KEGG modules associated with starvation-enriched genomes.

### Antibiotic resistance testing

Utilizing a broad array of antimicrobial compounds, we assess the antibiotic resistance of three nares polymicrobial communities from samples N3, N5, and N6, three saliva polymicrobial communities from samples S2, S5, and S8, and the three isolated strains of *Klebsiella* (N9-2-1, N11-KP-1, and S27-4-1). These were all SHI-medium communities before starvation. The selected nares and saliva communities did not contain any *Klebsiella* species by our 16S rRNA sequencing analysis. We also picked these three *K. pneumoniae* strains because *K. pneumoniae* is known to be more pathogenic than *K. aerogenes,* and they were cultured from three different individuals, representing both oral and nares strains. Resistance was assessed on brain heart infusion (BHI) supplemented with 5% Sheep’s Blood agar plates containing 250 μg/ml erythromycin, 50 μg/ml kanamycin, 35 μg/ml ciprofloxacin, 100 μg/ml ampicillin, 100 μg/ml streptomycin, 50 μg/ml rifampicin, 25 μg/ml nalidixic acid, 50 μg/ml vancomycin, antibiotic cocktail of *H. pylori* selective supplement (Oxoid; SR0147E), or sterile water. *H. pylori* selective supplement contained a mix of 2.5 μg/ml vancomycin, 1.25 μg/ml Trimethoprim, 1.25 μg/ml Cefsulodin and 1.24 μg/ml amphotericin. Overnight cultures of *Klebsiella* strains were grown in BHI broth aerobically at 37°C. Overnight cultures of nares and saliva communities were grown in SHI-media microaerophilically at 37°C. All cultures were serially diluted from 10^-1^ through 10^-4^, plated on each medium, and incubated overnight at 37°C microaerophilically. Growth on antibiotics was graded as either resistant (indistinguishable from growth without antibiotics), some resistance (reduced confluence), little resistance (few colonies present), or susceptible (no growth observed).

### Klebsiella pneumoniae polymicrobial community spike-in experiment

To experimentally test the enrichment of *Klebsiella* strains during starvation, three representative nares communities (N3, N5, and N6) and three saliva communities (S2, S5, and S8) that were completely susceptible to *H. pylori* selective supplement were spiked with *Klebsiella pneumoniae* N9-2-1 and incubated in nutrient-poor PBS (phosphate buffered saline). Out of the three antibiotic tested *K. pneumoniae* strains, we used *K. pneumoniae* N9-2-1 strain specifically in our experiment since this was the only strain cultured from day-120 starvation. We reasoned that those strains would have more selective advantage, and phenotype would be clearer to observe. An overnight culture of *K. pneumoniae* N9-2-1 was grown in BHI broth microaerophilically at 37°C. *K. pneumoniae* N9-2-1 was rinsed in PBS to remove nutritional carry-over before spike-in. Overnight cultures of nares and saliva communities were grown in SHI-medium microaerophilically at 37°C. Nares and saliva community cultures were normalized to an OD600 of 1.2 (approximately equivalent to 1.9 *10^6^ CFU/ml according to [35]), washed to remove residual growth medium, and resuspended in PBS. *K. pneumoniae* N9-2-1 was added to each community culture at approximately 0.1% (∼2,000 CFU/ml). This amount was estimated from our 16S rRNA sequencing analysis, which showed 0.1% to be the lower range of *Klebsiella* abundance in the community. Each *Klebsiella* spiked community was then aliquoted into 10 identical samples and incubated aerobically at 37°C. At 0 hours, 8 hours, 16 hours, 30 hours, 3 days, 12 days, 30 days, 60 days and 120 days after initiation, one set of samples were taken out of incubation, serially diluted through 10^-8^, and plated on both BHI supplemented with 5% sheep’s blood and BHI supplemented with 5% sheep’s blood and an *H. pylori* selective supplement (Oxoid; SR0147E). Growth in the absence of antibiotic selection indicated total culturable bacteria within the community, while growth in the presence of antibiotics indicated *Klebsiella*’s contribution to the culturable community.

For 16S rRNA gene sequencing studies, we used the same protocol as our antibiotic plating assay, except on day 30, we isolated genomic DNA and sent it for sequencing at the Zymo sequencing core (Table S10, S11, S12). For the nutrition-rich passaging experiment, instead of resuspending cultures in PBS, we used SHI-medium. Cultures were grown in 96-well plates at 250 μL of final volume. Every 24 hours, 50 μL of the incubated culture was passaged into a new 96-well plate containing 200 μL of fresh SHI-medium. After 30 days of passaging, the culture was pelleted, and genomic DNA was sent for 16S rRNA gene sequencing.

### Whole Genome Sequencing and Assembly

Out of the five nares and one saliva samples that had enriched *Klebsiella* after starvation, we were able to isolate two different *Klebsiella* strains from two individual nares samples (N9-2-1 and N11-KP-1) and one saliva sample (S27-4-1) (Table S9). Their genomic DNA was isolated (see above gDNA isolation method) and sent for whole genome sequencing at SeqCenter following their preparation recommendations. Based on the bacterial marker gene phylogeny (see next section), one of the *K. pneumoniae* strains (N11-KP-1) was identified as *K. variicola*.

All command line parameters used during genome assembly, binning, and analysis are described in full detail and can be found with our raw data on Zenodo (see Data Availability section). Briefly, each sample’s fastq files were quality trimmed with bbduk (https://sourceforge.net/projects/bbmap/) to remove adapters and drop low quality reads before quality filtering based on the recommendations of Minoche et al. 2011 [61], using illumina-utils [62]. Genomes were assembled using SPAdes [63]. Short reads were mapped back to each assembly and incorporated into anvi’o for manual binning [64]. Contigs were binned based on tetranucleotide frequency and evenness of coverage to remove any assemblies derived from contaminant DNA from the extraction and library preparation processes. Plasmid contigs identified as extrachromosomal at the time of sequencing (based on coverage disparity) were not included in the genome bin; other mobile elements with coverage patterns consistent with genomic integration were retained. plasmidSPAdes was also used to generate assemblies of any plasmids (https://academic.oup.com/bioinformatics/article/32/22/3380/2525610).

### Comparative genomic analysis

We used an iterative phylogenetic approach to identify a small set of genomes that adequately sampled the known diversity of *Klebsiella* relative to our new isolates. To accomplish this, we identified all *K. pneumoniae* genomes with readily accessible metadata on NCBI to create phylogenomic trees with GToTree [65]. Oversampled clades were pruned or dropped based on the relative distance to isolate genomes, and the analysis was repeated until the genomes currently displayed were obtained. While this pruning process does remove information and could alter the overall topology of the tree, information about body site or health association, if an evolutionarily conserved trait, should persevere. *K. variicola* genomes were obtained from a recent multi-hospital analysis [66] with detailed metadata on the isolation site and host health status. *K. variicola* At-22, the leafcutter ant symbiont, was also included to represent a known environmental genome (i.e., lineages not known to associate with mammals). Two *Klebsiella* genomes from the Baker *et al.* 2019 saliva starvation experiments (84-day saliva and PBS starved isolates) [34]. The final phylogenomic tree was generated with IQ-TREE2 [67] using the marker genes identified with GToTree.

All publicly available genomes were downloaded with ‘ncbi-genome-download’ and incorporated into anvi’o including annotation with KEGG, Pfam, NCBI COG [68–70]. A pangenome was constructed using ‘anvi-pan-genome’ with ‘--mcl-inflation 10’, which uses the Markov Clustering Algorithm to cluster amino acid alignments into “gene clusters”, putatively homologous groups of genes. The presence/absence of these gene clusters was then plotted for each genome and used to compare gene content similarity via a dendrogram. Average nucleotide identity (ANI) was also computed using pyANI [71] with the blast method, and the ANI values shown reflect the percentage over the alignable regions. The presence of metabolic modules, including predicted antibiotic resistance pathways, was estimated using ‘anvi-estimate-metabolism; based on KEGG pathways [72]. Antibiotic resistance was estimated with AMRFinderPlus using the *K. pneumoniae* model (Table S13, S14) [73].

## Data Availability

All relative abundance data are provided in the manuscript supplemental tables. Raw nucleic acid sequences and code used in this project are available on Zenodo. The raw data and code are also available at https://www.borlab.org/resources. Bacterial strains used in this paper will be provided upon request.

## Supplemental Figures

**Figure S1.**
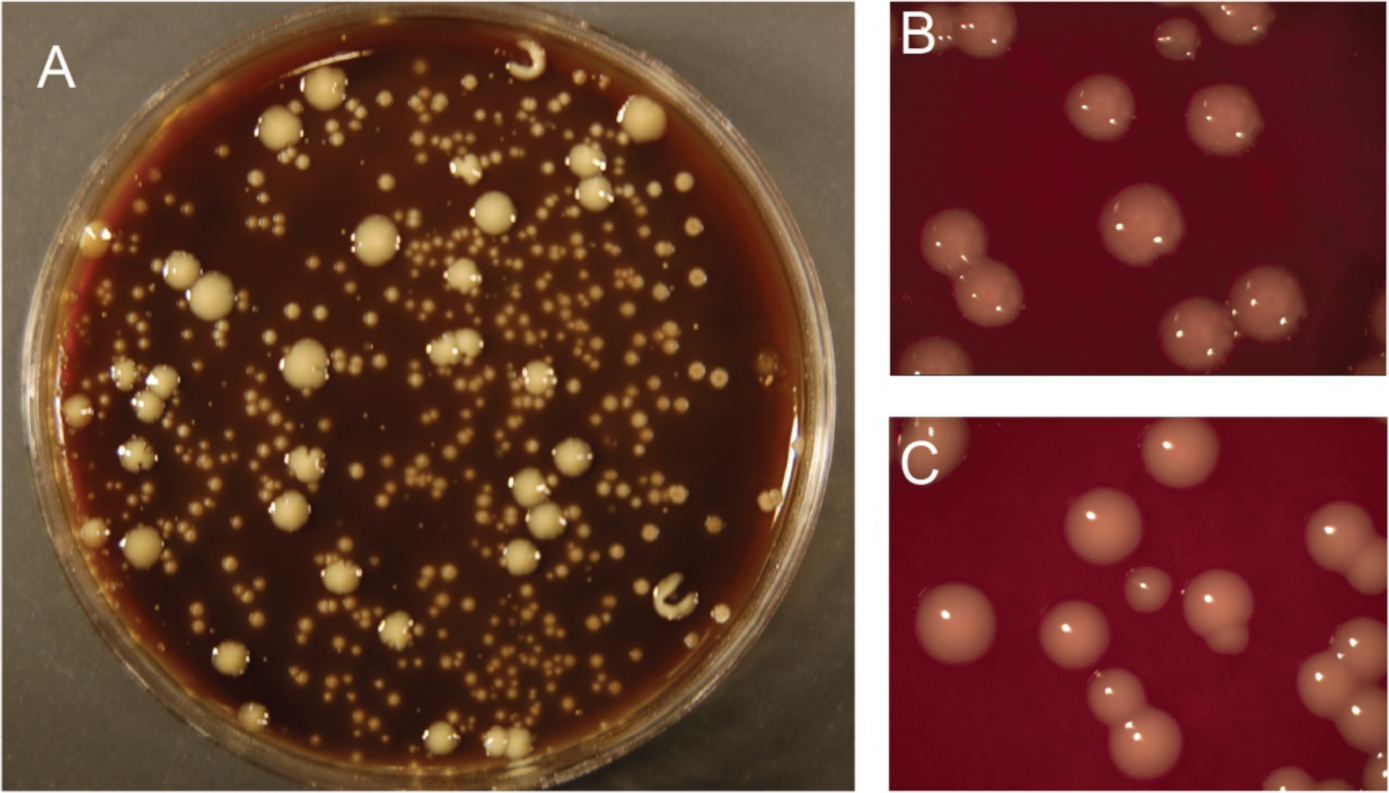
Extended starvation results in homogeneous colonies. (A) Saliva directly plated on blood BHI agar shows many colony sizes and types. (B-C) After day-30 and day-120 starvation, these colonies become homogeneous in size and morphology, many displaying mucoid phenotype. Numerous images were taken and only representatives are shown.

**Figure S2.**
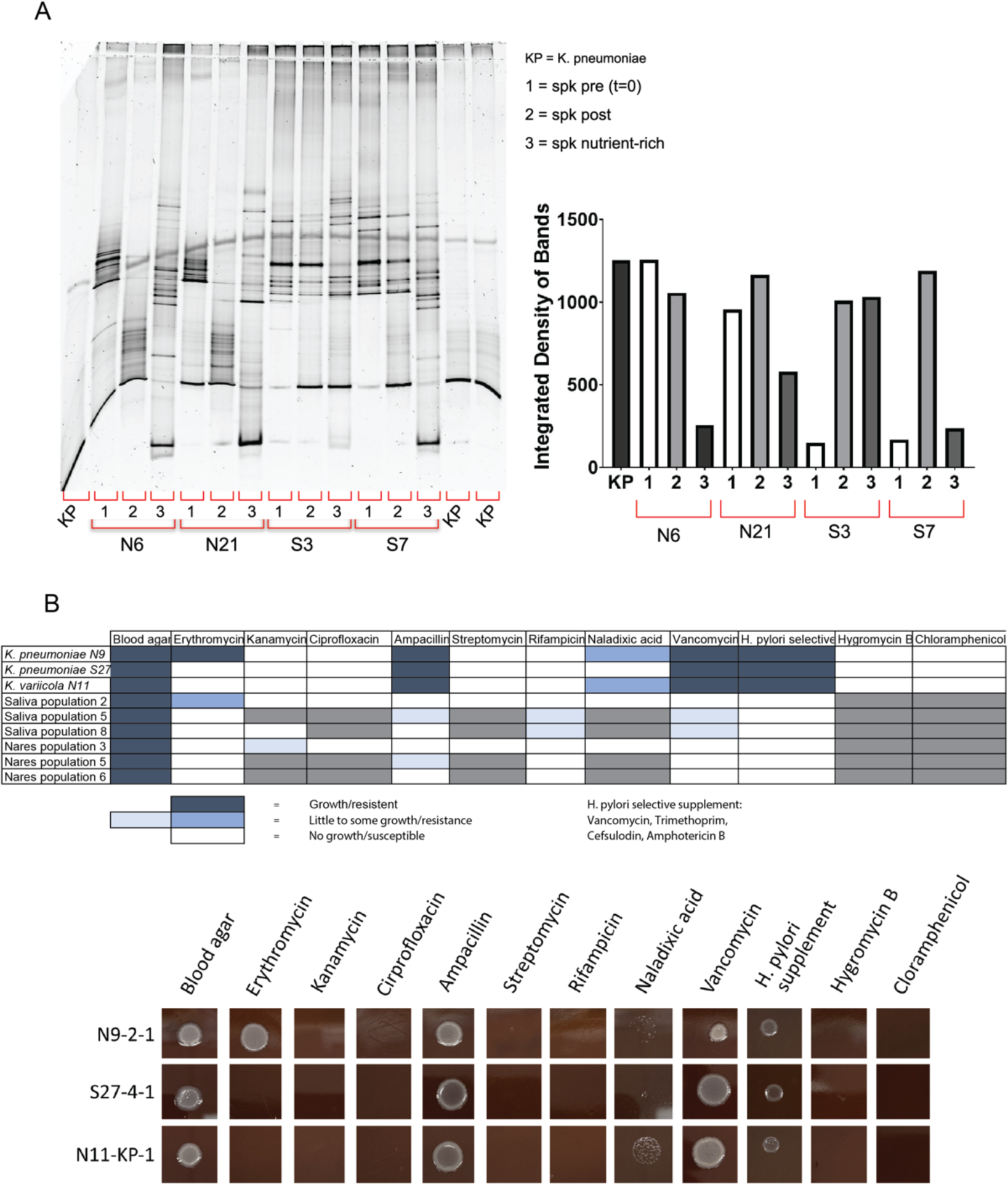
Characterization of isolated and cultured *Klebsiella pneumoniae* strains. (A) Artificial spiking of saliva and nares communities with *K. pneumoniae* followed by 30 day starvation resulted in increased amount of *K. pneumoniae.* After starvation, gDNA of the community was isolated and processed for DGGE. Samples were ran on large gels, and the bands were send for sequencing to identified the bacteria, as well as image was taken to quantify the band size and intensity. Total band gray area was calculated using ImageJ and plotted on the right. (B) Screening of isolated *Klebsiella* strains, nares communities, and saliva communities using a range of antibiotics identifies selective agents which can be used to isolate *Klebsiella* strains from a mixed culture. The table indicates the relative growth of each culture on the indicated antibiotic selection. Dark blue indicates robust confluent growth, lighter shades of blue indicate impaired growth, white indicates no visible growth, and grey indicates a combination which was not tested. Representative growth of the isolated *Klebsiella* strains on each tested antibiotic show strain variation in resistance profile.

**Figure S3.**
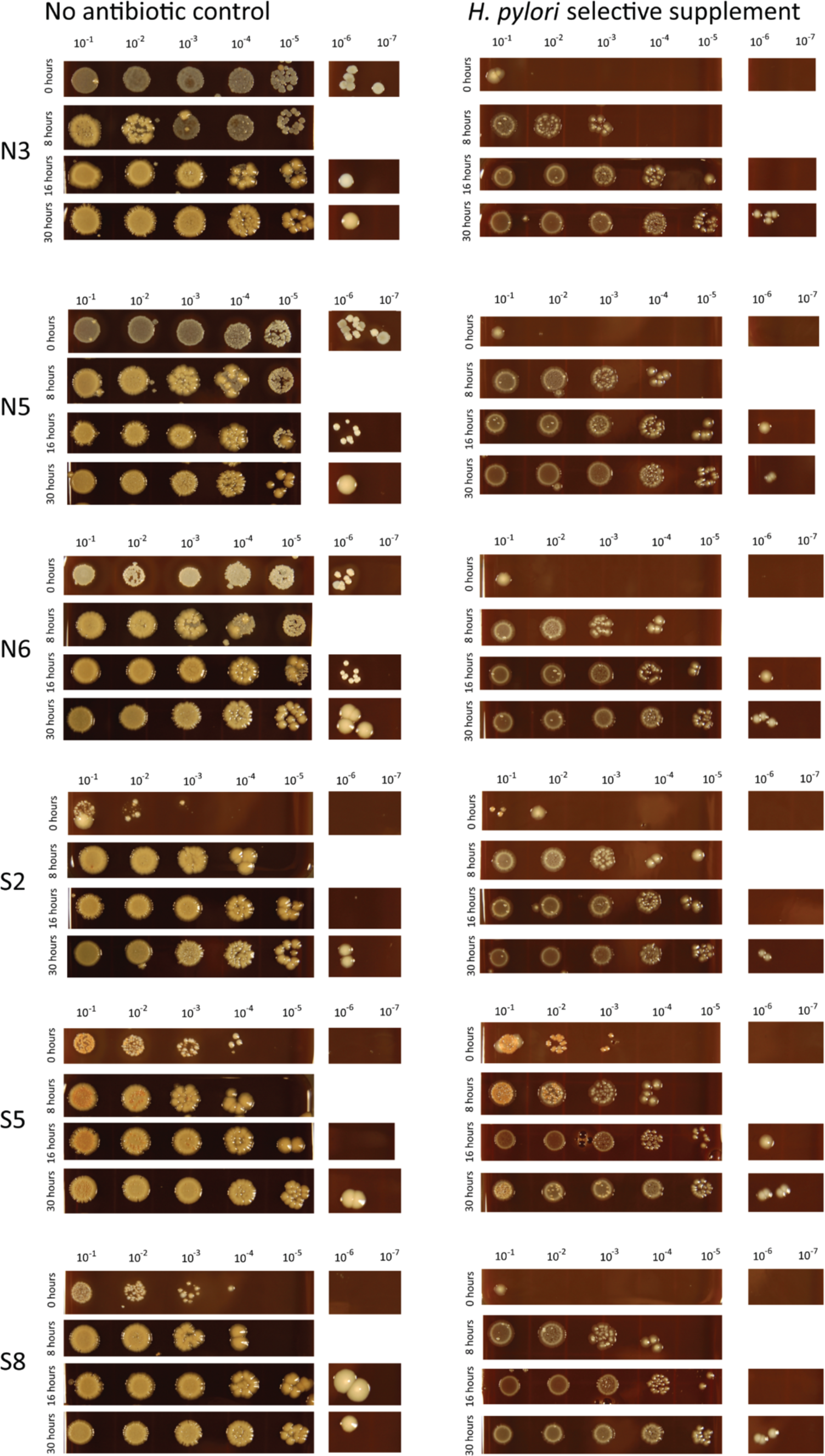
Longitudinal *K. pneumoniae* spiking and starvation experiment. Serial dilution of bacterial communities from the nares (N3, N5, N6) and saliva (S2, S5, S8) which have been inoculated with an oral *K. pneumoniae* (N9-2-1) strain demonstrate rapid domination by *K. pneumoniae* when incubated in nutrient poor PBS. At 0, 8, 16, and 30 hours after starvation bacterial communities were serially diluted from 10^-1^ to 10^-7^ and 20 μL of each dilution was spotted on non-selective BHI blood plates (shown on the left) and *Klebsiella* selective media (shown on the right). *K. pneumoniae* becomes the predominant culturable bacteria within nares communities after 30 hours of starvation and within saliva communities after 8 hours. All bacterial spots were performed in technical triplicate and only representatives are shown.

**Figure S4.**
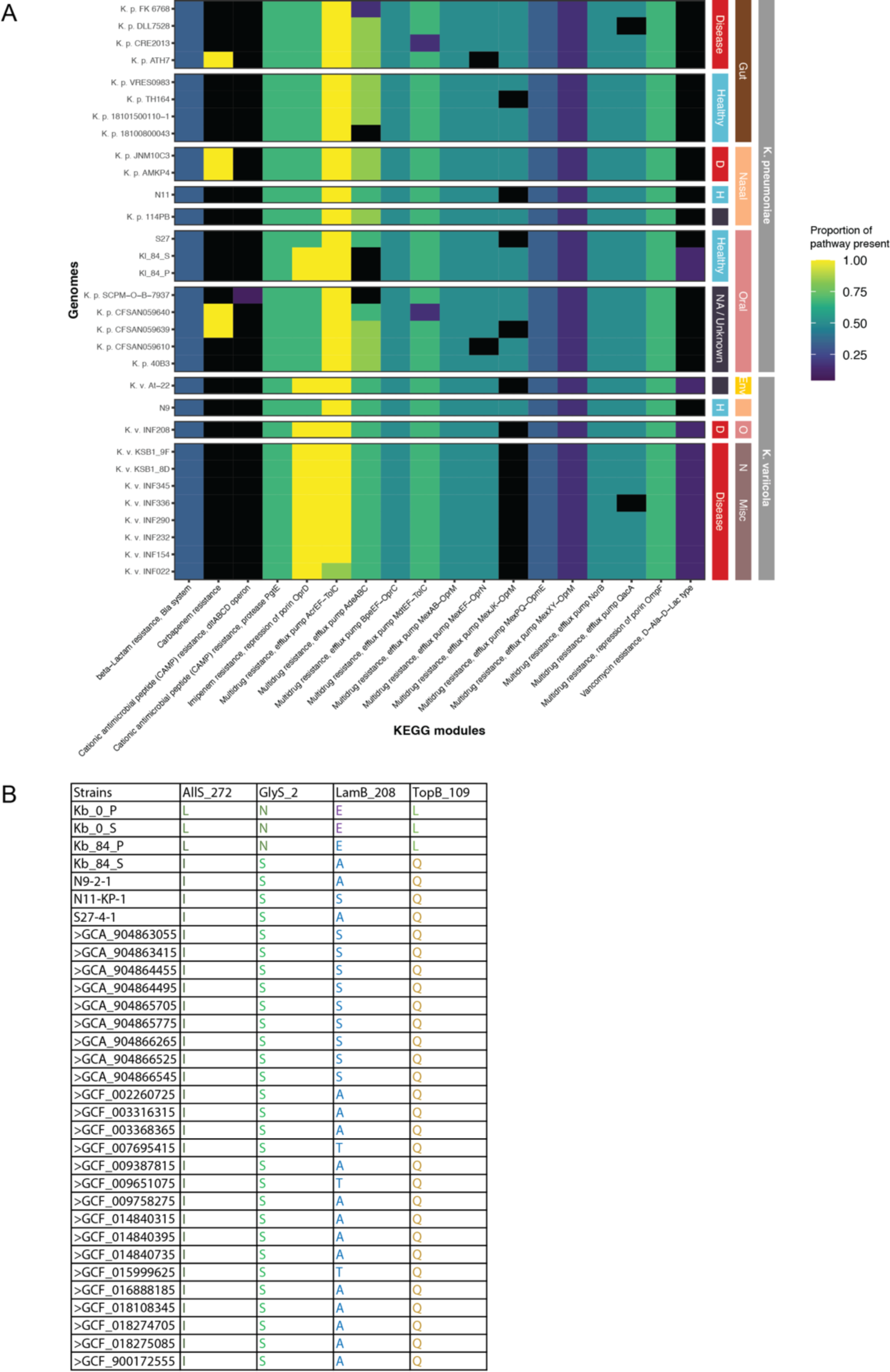
Genome analysis of isolated *Klebsiella* species from the starved saliva and nares samples. (A) Detection of KEGG metabolic pathways (columns) related to drug resistance, pathogenicity, and symbiosis across representative genomes (rows). Cells are colored by the proportion of each pathway’s genes present in each genome, i.e., 1 represents all genes in the pathway were detected. Genomes are arranged by isolation source and host health status (where relevant), demarcated with colored boxes on the right. (B) Amino acid residues found in each genome for the positions identified by Baker et al. 2019 as under selection during starvation.

**Figure S5.**
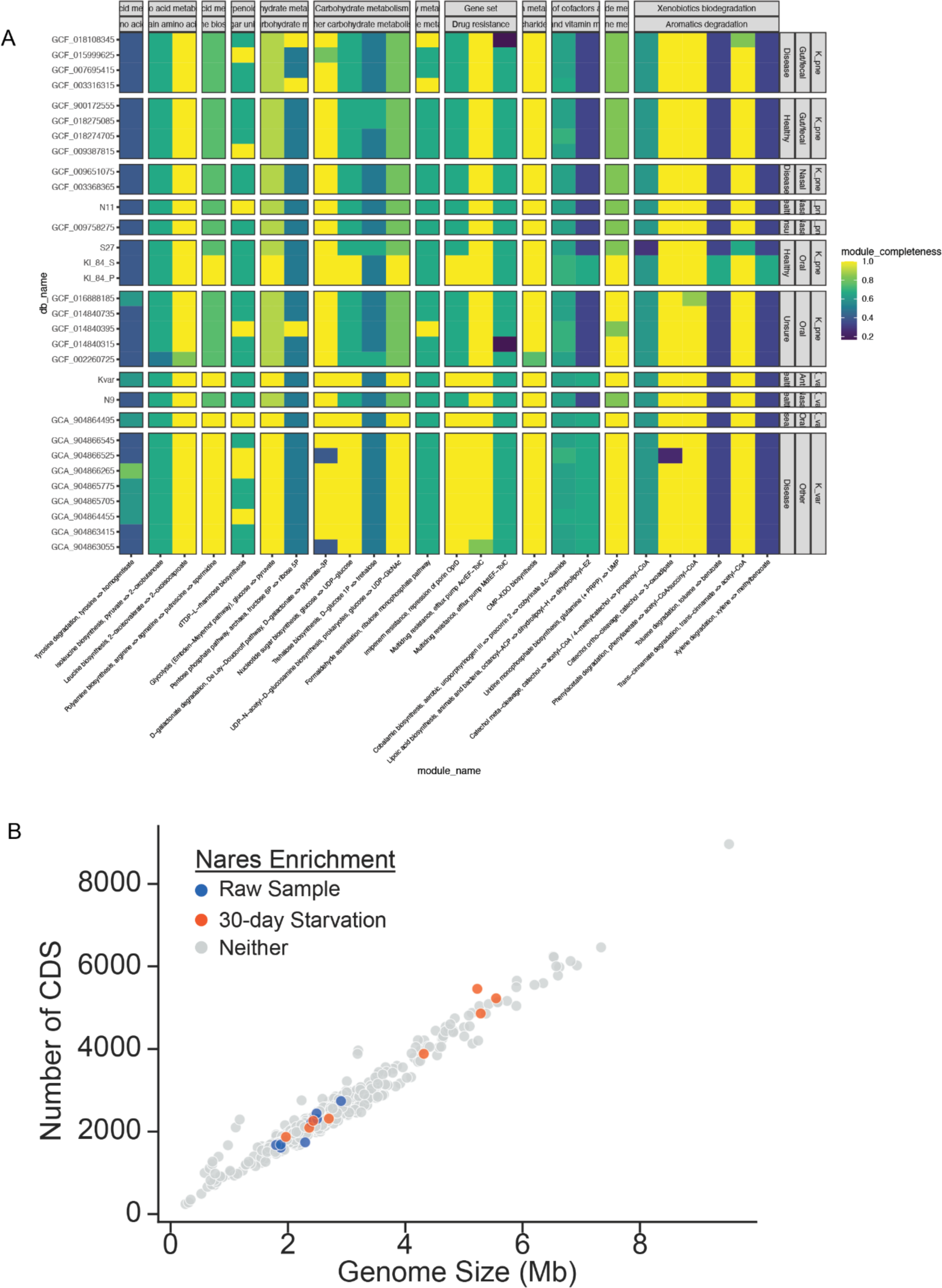
Global genome and genome size analysis of isolated *Klebsiella* strains. (A) Major metabolic pathways detected in representative *Klebsiella* genomes. Cells are colored by the proportion of each pathway’s genes present in each genome. Note that the figure shows only metabolic pathways with at least half (0.5) of the expected genes present. Genomes are arranged by isolation source and host health status (where relevant). (B) Genome of bacterial species enriched in raw (blue) or day-30 starved (orange) samples were obtained from the eHOMD. The size of the genomes were plotted against all other oral and nasal genomes (gray) from the eHOMD.

